# Habituation drives elevated antimicrobial resistance, functional, and compositional shifts in the black bear gut microbiome

**DOI:** 10.64898/2026.09.25.754533

**Authors:** Summer E Vance, Jillian Adkins, Dylan Maghini, Robert W Crawford, Ami S Bhatt, Elizabeth Hadly

## Abstract

Wildlife habituation across the globe poses threats to human and animal safety and alters regular ecological function, yet the wildlife, environmental, and public health ramifications are poorly characterized. The gut microbiome is a valuable proxy for these factors as it is affected by anthropogenic exposure, influences host health, and may contribute to the environmental spread of virulence factors. Despite their importance, wildlife gut microbiomes are largely uncharacterized, both in composition and function. We characterized American black bear (*Ursus americanus*) gut microbiomes across a spectrum of lifestyles including wild (conflict-free), habituated (conflict-prone), and captive individuals. 16S rRNA gene sequencing was used for compositional analyses and whole genome short-read sequencing (WGS) was performed on a subset of bears to investigate microbiome function and virulence factors. We found that habituated, wild, and captive bears have unique gut microbiome compositions that are indicative of their respective lifestyles. In particular, captive bears are enriched in carnivory-associated gut microbiomes and habituated bears in inflammatory gut microbiomes. WGS reveals wild bears have increased fungi and soil-associated bacteria, as well as higher proportions of unclassified reads. Habituated bears have significantly increased levels of antimicrobial resistance genes, and captive bears are enriched in enterotoxins. We demonstrate that consequences of wildlife habituation corroborate a One Health framework, showing the interconnectedness of human, animal, plant, and environmental health.

**Graphical Abstract:** 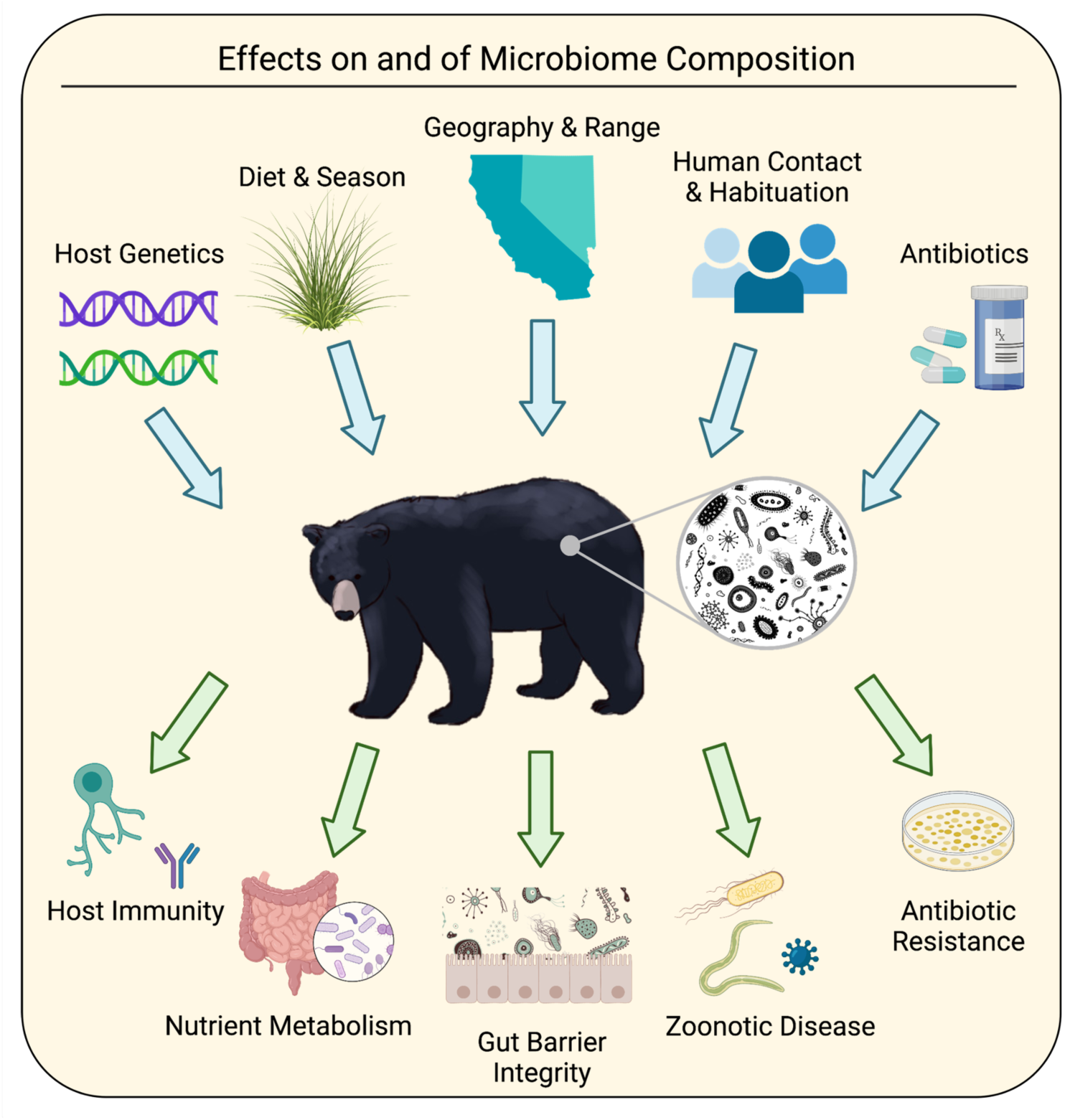

## INTRODUCTION

In a world of rapid urbanization, wildlife is increasingly exposed to humans, urban spaces, and human-sourced nutrients, regularly resulting in habituation. Wildlife habituation occurs with repeated exposure to the same stimuli, in this case human contact, eventually resulting in wild animals no longer responding to that stimulus (Mazur). Wildlife habituation is characterized by a lack of fear of humans and vehicles, and is often associated with conditioning, where wildlife seeks and consumes human-sourced nutrients such as human and pet food, trash, toiletries, compost, etc. (Mazur). Habituation and conditioning result in increased incidence of human-wildlife conflict and negative outcomes. Because habituation also alters wildlife behavior (Ritzel and Gallo; Wilmers et al.; Goumas et al.), it exposes wildlife to trash, pharmaceuticals, toxins and other anthropogenic pollutants that may negatively influence health (Larson et al.; Murray et al.). In long-lived animals with prolonged maternal care, such as bears, habituated behaviors are passed down to offspring, perpetuating conflict across seasons and years (Schell et al.; Shimozuru et al.). Habituated animals are also at higher risk of human-caused mortality, including being hit by vehicles, ingesting poison or toxins, and management euthanasia (Gantchoff et al.).

Black bears (*Ursus americanus*) have a long history of problematic habituation in the United States, resulting in several established wildlife management and “rewilding” programs particularly in national parks (Mazur). Black bears are omnivorous generalists whose opportunistic nature and high intelligence make them both susceptible to and successful in habituation. Increased habituation is linked to increased boldness and aggression, which may pose serious threats to people and property. Conflict-prone bears are often lethally removed (Mazur). While management programs such as bear-proofing trash containers have successfully reduced property damage, isotopic evidence indicates that black bears in Yosemite National Park still consume human-sourced food at levels comparable to the early 1900s (Hopkins et al.). In addition, management programs are largely confined to highly managed protected areas, while black bear ranges are widespread across North America (Kirby et al.).

Thus, habituated, food-subsidized black bears persist, yet the sublethal impacts of habituation on bear health have not been well-documented.

Here, we use the bear gut microbiome as a non-invasive measurement of physiological health, which is prohibitively difficult to measure directly for wildlife. Gut microbiome composition is a crucial indicator of host health and has been linked to disease states, physiology, immunology, and neurological function in humans and model organisms (Gebrayel et al.).

Wildlife microbiomes are largely uncharacterized both in community structure and function, though these hosts are equally influenced by their microbial inhabitants (Lagerstrom et al.). Importantly, microbiomes also enact effects outside of the host, and thus doubles as environmental health proxies. Pathogenic factors in the microbiome such as certain bacterial species, viruses, antimicrobial resistance genes (AMR), and toxins (Trinh et al.) impact the host and are excreted in feces to the environment, where they may be transferred to other wildlife, livestock, pets, and humans (van Bruggen et al.). Thus, understanding wildlife microbiome composition is an important component of One Health, as it integrates wildlife, environmental, and public health factors (van Bruggen et al.; Trinh et al.).

A growing body of work examines the effects of anthropogenic stressors on wildlife microbiomes (Fackelmann et al.) including captivity (Schwab et al.; Moustafa et al.), climate warming (Greenspan et al.; Hernández-Gómez), habitat fragmentation (Amato et al.), and land use change (San Juan et al.; Watson et al.; Heni et al.). Decreased microbiome diversity is associated with captivity in many mammalian species (Moustafa et al.; Clayton et al.; Jin et al.; Borbón-García et al.). Much of the work on captive animal microbiomes also documents “humanization” of the gut microbiome, which identifies microbe-sharing between animals and humans (Dillard et al.; Clayton et al.). The effects of urbanization and anthropogenic food consumption on wildlife gut microbiota are less well-documented, and the few existing studies have contradictory findings. Consumption of anthropogenic food (with land urbanization used as a proxy) has variably been shown to decrease microbiome diversity (Sierra J Gillman et al.), not affect diversity (W. Lee et al.; Xia et al.), or increase microbiome diversity (Sugden et al.; Teyssier et al.; Phillips et al.). Overall, changes in gut microbiomes across gradients of anthropogenic disturbance and human-sourced food consumption seem to be highly species-specific (Heni et al.). Additionally, there is a dearth of research on wildlife microbiome pathogenicity and its role in environmental health.

Here, we investigated the gut microbiome composition of black bears across an anthropogenic exposure gradient, including wild (conflict-free), habituated (conflict-prone), and captive individuals. We performed 16S rRNA gene sequencing on 65 fecal samples with additional short-read Illumina sequencing on a subset of 17 individuals to interrogate microbiome functionality and pathogenicity. We used 16S rRNA gene to characterize microbiota composition, and whole genome short-read sequencing (WGS) to further identify antibiotic resistance and toxin-associated genes, and functional profiles. The vast majority of wildlife microbiome studies have used only 16S rRNA gene sequencing (Combrink et al.), which cannot adequately tease apart microbes at low taxonomic levels, detect microbes at low abundance, or profile genes (Durazzi et al.). Deeper WGS methods allow us to more thoroughly examine microbiome composition and function. In WGS work that has been done on wildlife microbiomes, researchers find vast amounts of unclassified biological and functional diversity (Bohra et al.; Cabral et al.; C. Song et al.; Jin et al.). Our work is unprecedented in its inclusion of three distinct wildlife lifestyle cohorts and the power of WGS to address multiple virulence factors as well as microbiome function.

## METHODS

### Black bear fecal sample collection

Fecal samples were collected from representatives of captive, wild, and habituated experimental groups of *Ursus americanus* from California and western Nevada. Fecal samples were collected from representatives of captive, wild, and habituated experimental groups of *Ursus americanus* from California and western Nevada. These cohorts represent distinct lifestyles with differential exposure to humans and anthropogenic food and pollutants. Briefly, wild bears have minimal or no contact with humans or urban spaces and source food from natural habitats, typically consuming grasses and vegetation in the spring and early summer, then preferentially consuming nuts and berries later in the summer season, with opportunistic meat consumption year-round. Habituated bears live near urban areas and have increased contact with humans and anthropogenic pollutants. The bears included in this study were identified as “conflict bears” due to their presence in urban spaces, where they typically consume human waste from dumpsters and show a reduced fear of humans. Captive individuals were housed in either zoos or rehabilitation centers where they interacted with caretakers, were fed standardized diets, and had access to medical treatment.

Sample collection occurred from January 2017 through November 2018. Fecal collection kits containing a sterile 250 mL conical tube, sterile tongue depressor, bleach wipe, and collection instructions were distributed to select captive facilities within California as well as California Department of Fish and Wildlife (CDFW) and Nevada Department of Wildlife (NDOW) biologists. After collection, samples were stored at -20°C until shipment to the CDFW Wildlife Forensics Lab where they were then stored at -80°C.

### 16S metagenomic sequencing

DNA was extracted from each fecal sample using the QIAamp Fast Stool Mini kit (Qiagen, Inc) following manufacturer’s instructions with the addition of a homogenization step using pre-filled DNase free 2 mL tubes containing 0.5mm glass beads and incubation of samples with a 15 mg mL^-1^ lysozyme solution at 37°C for 30 minutes with vortexing every 10 minutes. Homogenization of samples was done using Analytik Jena SpeedMill PLUS for three consecutive cycles of 60 seconds with 60 second cooling between cycles. Extracted DNA was eluted in 100 µL of dH_2_0. DNA yield was quantified after each extraction and quality assessed using a NanoDrop spectrophotometer (ThermoFischer, Inc) and Quantus fluorometer (Promega, Inc). DNA was amplified using the KAPPA HotStart Ready Mix with barcoded primers targeting the V3 and V4 hypervariable regions of the 16s rRNA gene selected from Klindworth et al. (Klindworth et al.) using the following thermocycling conditions: 95°C for 3:00; 30 cycles at (95°C 0:30, 55°C 0:30, 72°C 0:30), 72°C 5:00; 10°C hold. Dual indexing primers were added to the first-step PCR product using the Nextera XT Index kit (24 or 96 indices). The pooled library was sequenced using a 600 cycle Miseq reagent Kit v3 according to Illumina’s 16s Metagenomic Sequencing Library Preparation guide.

Resulting sequences were analyzed using the QIIME2 (v2022.11.1) software package (Bolyen et al.). Demultiplexed sequences were filtered for quality, trimmed, and denoised using DADA2 (v 2022.11.2; Callahan, McMurdie, Rosen, et al.). We opted to use amplicon sequence variants (ASV) rather than operational taxonomic units (OTU) to decrease reliance on existing databases and increase resolution (Callahan, McMurdie, and Holmes). ASV taxonomy was assigned using the q2-feature-classifier (Bokulich et al.) against the Greengenes reference database (DeSantis et al.). Differential abundances of families and phyla were determined using DESeq2 (v3.17; Love et al.).

### Short-read metagenomic sequencing

Samples for short-read sequencing were chosen to minimize seasonal diet change. DNA was extracted from frozen scat using the Qiagen QIAamp PowerFecal DNA Kit (QIAGEN Cat. No. 12830) using the manufacturer’s recommended protocol. DNA concentration was measured with Qubit Fluorometric Quantitation (DS DNA High-Sensitivity Kit, ThermoFisher Cat. No. Q32851). DNA purity was measured with a NanoDrop 2000 Spectrophotometer (ThermoFisher Cat. No. ND2000). Sequencing libraries were prepared using the Nextera Flex DNA Library Prep Kit (now listed as “Illumina DNA Prep,” Illumina Cat. No. 20018704). Library concentrations were measured using Qubit Fluorometric Quantitation and library size distributions were analyzed with the Bioanalyzer 2100 (Agilent G2939BA). Libraries were multiplexed and 150 bp paired-end reads were generated on the HiSeq 4000 platform (Illumina).

Short-read sequencing data was demultiplexed by Novogene. Demultiplexed data were run through a publicly available preprocessing pipeline (https://github.com/bhattlab/bhattlab_workflows/blob/master/preprocessing/preprocessing.snakefile, commit version ec0af84 as of Dec 19, 2021). TrimGalore (https://github.com/FelixKrueger/TrimGalore, v0.6.5) is used to trim reads with a minimum quality score of 30 and minimum read length of 60. Trimmed reads are then deduplicated using *htstream* SuperDeduper (v1.2.0) with default parameters (Petersen et al.). Because no satisfactory black bear genome exists, host reads were removed by filtering out all Metazoa hits. Assembly was run via a publicly available pipeline (https://github.com/bhattlab/bhattlab_workflows/blob/master/assembly/assembly.snakefile, commit version ec0af84 as of Dec 19, 2021) using the assembler MEGAHIT (v1.0) (Li et al.).

Results are evaluated using QUAST (v4.6) (Gurevich et al.). Classification of assemblies was run using a publicly available pipeline (https://github.com/bhattlab/kraken2_classification/blob/master/Snakefile, commit verison dd2928e as of Dec 15, 2021). The pipeline utilizes Kraken (v2.0.9-beta) (Wood et al.) with default parameters and a comprehensive custom reference database containing all bacterial and archaeal genomes in GenBank as of January 2020 (Clark et al.). Bracken (v2.2.0) is then used to re-estimate abundance at each taxonomic rank (Lu et al.). Reads classified as *Chordata* are removed from the final output. Differential abundances of genera were determined using DESeq2 (v3.17) (Love et al.).

### Virulence factors analysis

ShortBRED (https://github.com/biobakery/shortbred.git, commit version 22596ef as of Mar 23, 2021) was employed to profile virulence factors using default settings, where a hit is recorded if the marker-read alignment has 95% or more identity and is at least as long as the shortest marker length or 95% of the read length (Kaminski et al.). Reads are aligned against the virulence factor database (VFDB) (Liu et al.). We then further filtered hits for E-values < 0.001.

### Antimicrobial-resistance gene analysis

Resistome data was generated using the Resistance Gene Identifier (RGI v5.2.1), which uses homology and SNP models to match assemblies to the Comprehensive Antibiotic Resistance Database (CARD v3.1.4) (Alcock et al.). Open reading frames were identified using a gene prediction algorithm performed by Prodigal (v2.6.3) (Hyatt et al.). Results were filtered to 80%, as well as 100% alignment to identify.

### Toxin-associated gene analysis

PathoFact (v1.0) was used to predict toxin genes in the short-read assemblies (De Nies et al.). The *hmmsearch* function of HMMER3 (v3.2.1) matches query assemblies to the toxin HMM database, which is composed of SwissProt, KEGG, PFAM, and TIGR toxin-associated gene sequences (Finn et al.). These results are combined with SignalP (v5.0) to classify the hits as associated with secreted or non-secreted toxins (Almagro Armenteros et al.). We further grouped the toxin library of 465 genes into enterotoxins, hemolysins and cytolytic toxins, other toxins, efflux pumps and transporters, toxin-antitoxin system, toxin associated enzymes, toxin associated domains, and other toxin associated genes.

### CAZyme composition

Run_dbcan (V4, https://github.com/linnabrown/run_dbcan.git, commit version 42af6e7 as of Jan 24, 2024) was used to analyze CAZyme profiles. Run_dbcan identifies putative carbohydrate-active enzymes using diamond, hmmer, and DataBase for automated Carbohydrate-active enzyme ANnotation (dbCAN). We then filtered for CAZymes that were detected by all three tools.

## RESULTS

Single fecal samples were collected from 65 bears spanning an anthropogenic exposure gradient, including wild bears (n = 22), habituated bears (n = 22), and captive bears (n = 21) (Fig. 1, Table S1). Wild bear samples were collected non-invasively and opportunistically from Los Angeles National Forest, whereas habituated bear samples were collected during captures of conflict-prone individuals in the southern and eastern Lake Tahoe regions, and captive bear samples were collected opportunistically from care facilities. To measure microbiome composition, we performed 16S rRNA sequencing on all samples and generated a median of 55,448 (range 3,956 -101,487) amplicon sequence variants (ASV) per sample. To enable deeper functional characterization of microbiomes, we performed 2 x 150 base pair paired-end shotgun sequencing on a subset of bears from each group (wild, n = 7; habituated, n = 5; captive, n = 5), generating a median of 46,374,645 (range 68,199,064 -21,555,207) reads per sample with a median of 28,753,559 (range 14,393,123 -43,520,538) reads remaining after quality control. Bears were selected for shotgun sequencing based on DNA extraction quality and month of collection to minimize seasonal bias.

**Figure 1.**
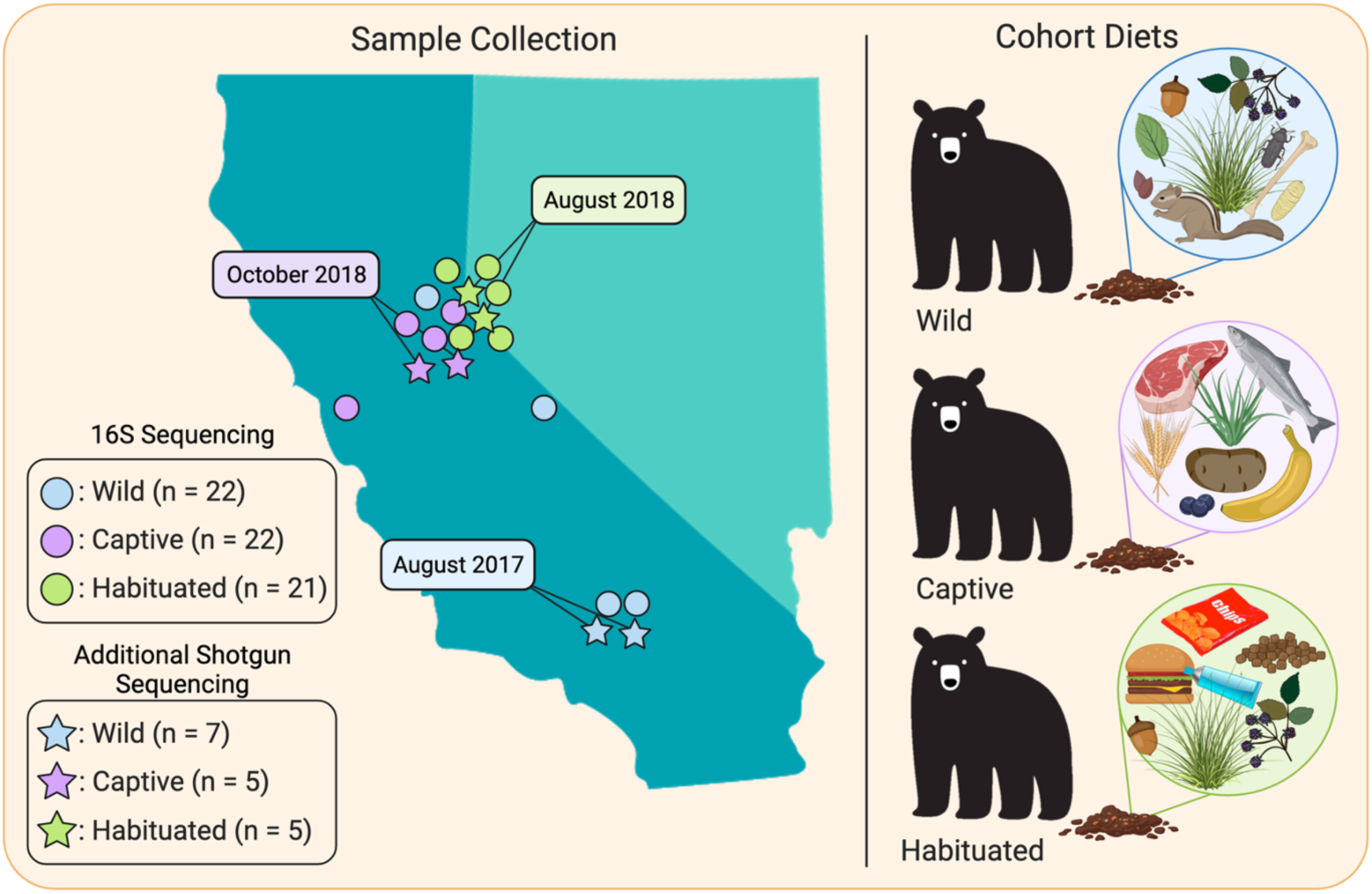
Study design and sample collection. Points of sample collection across California and Nevada between January 2017 and November 2018. Wild bears are shown in blue, captive in purple, and habituated in green. Circles represent samples used for 16S sequencing only. Stars represent samples used for 16S and shotgun sequencing. Diets differ between cohorts.

### Microbiome composition shifts across lifestyles

First, we investigated the overall differences in microbial composition between bear populations with our 16S rRNA sequencing. Shannon diversity, a metric that represents the within-sample taxonomic diversity of a microbiome, was not significantly different between groups (Fig. 2a).

**Figure 2.**
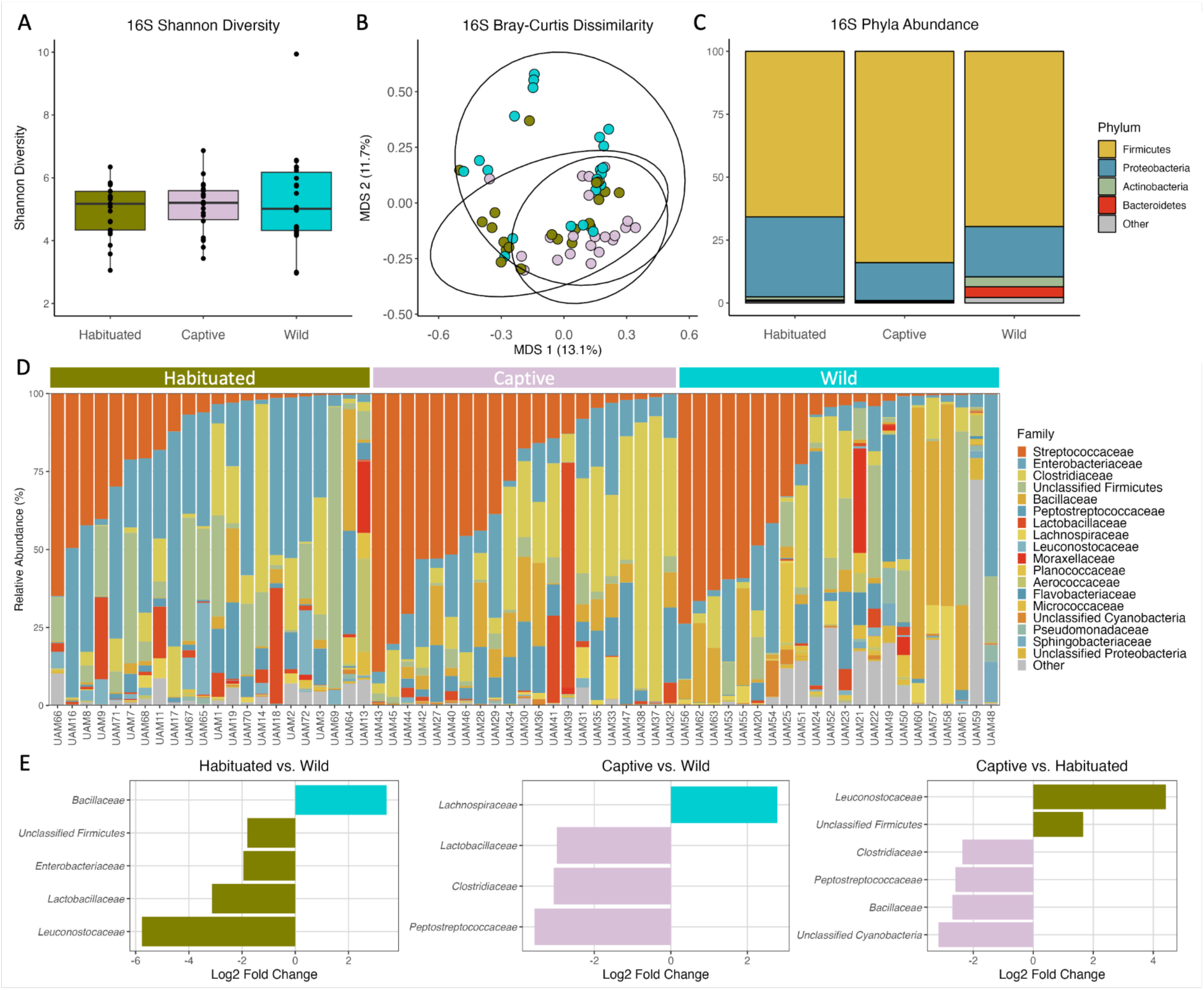
16S composition. A. Shannon diversity by cohort. B. Bray-Curtis dissimilarity matrix colored by cohort. C. Phyla abundance by cohort. D. Family relative abundance by individual, grouped by cohort. E. Pairwise significantly different bacterial families.

Bray-Curtis dissimilarity, a measure of compositional differences between microbiomes, indicates that bears from each group do have microbial taxonomic profiles that cluster together (p = 0.001; permanova test). Pairwise differences between cohorts all remain significant even after Benjamini-Hochberg correction (habituated vs captive, p = 0.024; habituated vs wild, p = 0.018; captive vs wild, p = 0.006). The first and second axes of variation explain 13.1% and 11.7% respectively of the overall compositional variance (Fig. 2b). These results imply that while the overall within-sample diversity is similar across bear populations regardless of lifestyle, there may be taxonomic differences between groups that differentiate their microbiomes at the community level.

To specifically identify the microbial taxa that drive these differences, we calculated the abundance of microbes at the phylum and family levels. At the phylum level, Firmicutes are the most abundant across all cohorts, followed by Proteobacteria and lower abundances of Actinobacteria and Bacteroidetes (Fig. 2c). Habituated bears have significantly increased Proteobacteria compared to the other cohorts (mean abundance = 45.10%; habituated vs captive, p = 0.029; habituated vs wild, p = 0.043; anova with post-hoc Tukey HSD). At the family level across all bears, *Streptococcaceae* is the most abundant, followed by *Enterobacteriaceae*, *Clostridiaceae,* and *Bacillaceae*, although composition varies between individuals within each group (Fig. 2d). Several families are significantly differentially abundant between groups (Fig. 2e). Of particular interest, *Peptostreptococcaceae* and *Clostridiaceae* are enriched in captive bears relative to the other cohorts. *Peptostreptococcaceae* has been previously observed in carnivorous mammals (De Jonge et al.), which is consistent with the meat-enriched diet of captive bears. Similarly, *Clostridiaceae,* most often a commensal gut inhabitant despite containing several notorious pathogenic species, has also been described at very high abundance in the carnivorous polar bear gut microbiome (Watson et al.). Habituated bear gut microbiomes are enriched in *Enterobacteriaceae,* encompassing many taxa associated with inflammation (Moreira De Gouveia et al.), potentially reflecting the exposure of habituated bears to anthropogenic-associated pollutants.

These results are largely recapitulated in the subset of bears profiled with shotgun sequencing (Fig. S1 and S2); however, shotgun sequencing reveals a higher abundance of fungi and soil-associated bacteria in wild bears relative to captive and habituated bears. Further, a larger percentage of reads are unclassified in wild bears relative to captive and habituated bears, perhaps reflecting undescribed microbial novelty in the wild bear population. These results demonstrate that bear microbiomes reflect lifestyle differences across human-associated habitats, resulting in differences in the abundance of known dietary-associated taxa as well as taxa whose niches in wildlife microbiomes are poorly understood.

### Antimicrobial resistance gene prevalence is increased in habituated bears

Antimicrobial resistance (AMR) is a fast-growing threat to medical treatment of humans and animals worldwide. Though AMR is typically described in humans and livestock, investigating antimicrobial resistance in wildlife microbiomes can serve both as a potential marker for habituation and also to explore underappreciated reservoirs of AMR (Lagerstrom and Hadly). Using whole metagenome shotgun sequencing as input for antimicrobial profiling, we were able to classify AMR genes in every bear sample (Fig. 3a); this is expected since many microbes naturally code for AMR. However, we observe increased AMR abundance in captive bears (p = 0.032) relative to wild bears, suggesting an increased exposure to antimicrobials perhaps through direct treatment with antibiotics or consumption of antibiotic-treated food. Habituated bears also harbor significantly higher AMR abundance than wild bears (p = 9.6e-5) and appear more similar to captive bears, reflecting a strong effect of anthropogenically-sourced food.

**Figure 3.**
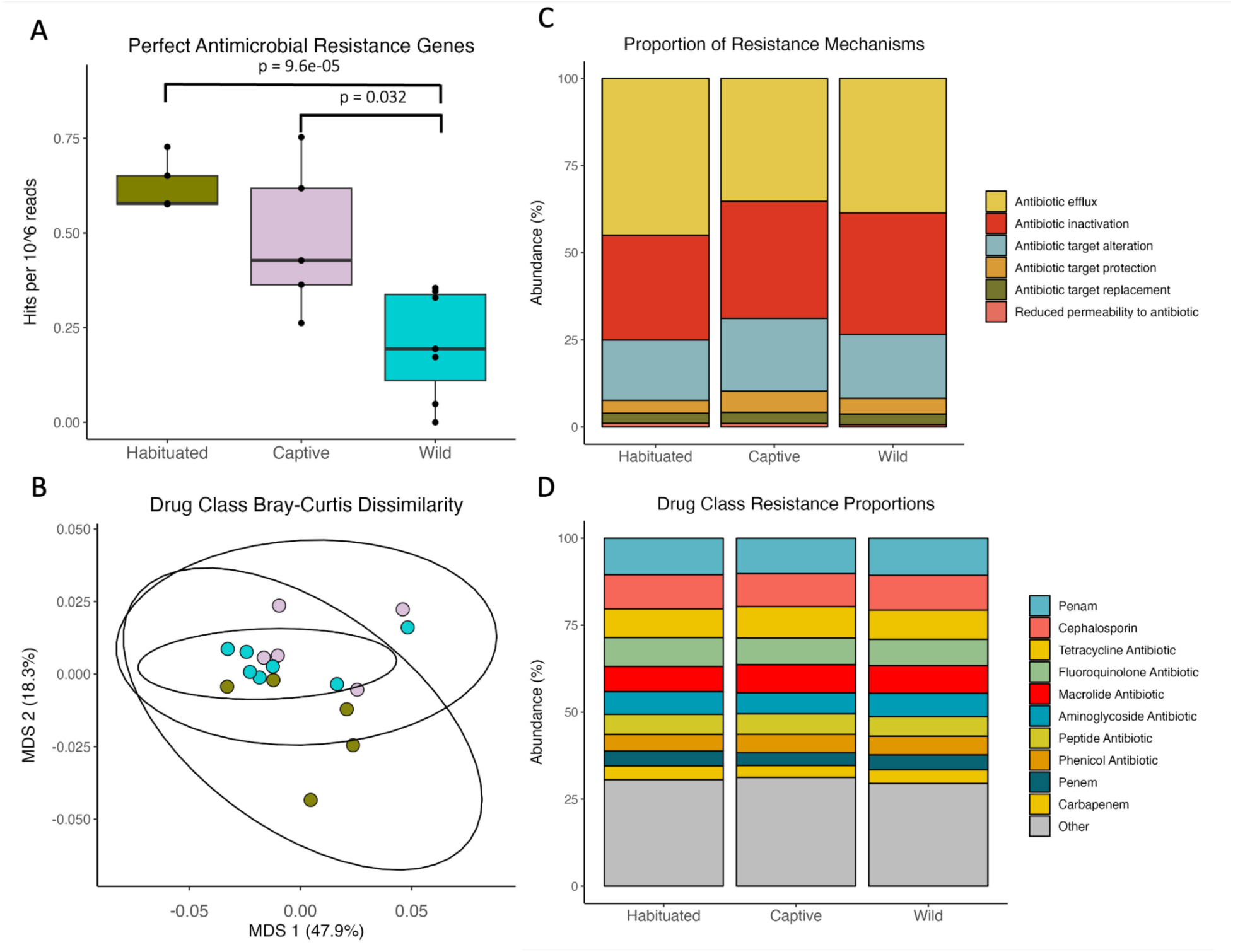
Antibiotic resistance gene profiles. A. Antimicrobial resistance genes per cohort normalized by read depth. B. Bray-Curtis dissimilarity MDS using antibiotic resistance gene profiles. C. Proportion of antibiotic resistance mechanisms by cohort. D. Proportion of drug class resistance by cohort.

To better understand the differences observed between groups, we classified the resistance mechanisms and targeted drug classes of each AMR gene, which both showed similar proportions across the cohorts (Fig. 3c and d). This suggests that captive and particularly habituated bears have increased AMR loads that are not driven by an increase in a specific AMR class, but rather a relatively equal increase across all AMR classes.

### Toxins, particularly enterotoxins, are increased in captive bears

Toxins are naturally produced by microbes and, when pathogenic, can be extremely detrimental to host health. In addition, gut microbe toxins are disseminated into the environment via feces. Although some wildlife microbiomes have been studied for the presence of specific toxins (e.g. shiga toxin; Persad and LeJeune), to our knowledge no wildlife microbiomes have undergone untargeted toxin profiling. As such, the relationship between lifestyle and toxin burden, and the potential of wildlife to serve as toxin reservoirs are poorly understood. Toxin gene profiles clustered by cohort in a Bray-Curtis dissimilarity matrix, with primary and secondary axes of variation describing 36.4% and 24.7% of the variance, respectively (Fig. 4c). Captive bears have significantly more secreted toxins than wild and habituated bears (p = 0.009, p = 0.021 respectively) (Fig. 4a), suggesting their diet and exposure to anthropogenic pollutants and/or environmental stress may make them more susceptible to colonization by toxin-producing microbes. Captive bears have significantly increased enterotoxin loads compared to wild and habituated bears (p = 0.008, p = 0.008) (Fig. 4b). Overall, toxin profiles show high within-group similarity (Fig. 4d). At the gene level, there were many significantly differentially abundant toxin-associated genes between groups (Fig. S3). Notably, we did not identify common toxins of concern, such as Shiga, cholera, *Clostridium difficile*, *Clostridium perfringens,* or *Clostridium botulinum* toxins, in any sample. Our results suggest that targeted toxin profiling overlooks a large amount of toxin diversity in wildlife microbiomes.

**Figure 4.**
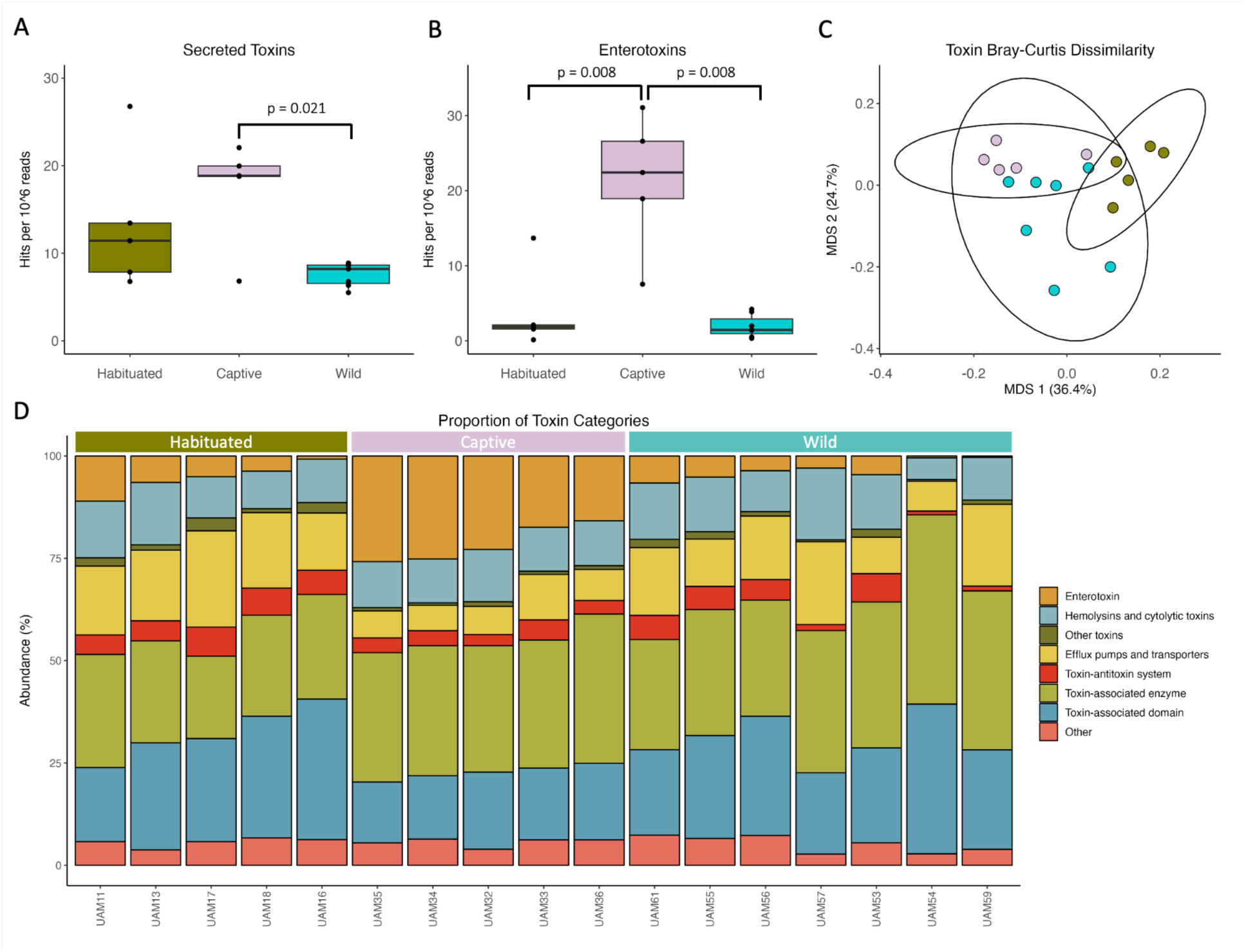
Toxin-associated gene profiles. A. Number of secreted toxin-associated genes per 1 million reads. Captive is significantly higher than wild (p=0.021). B. Enterotoxins are significantly increased in captive bears (p=0.008 against both habituated and wild cohorts). C. Bray-Curtis dissimilarity MDS using toxin profiles. D. Abundance of toxin-associated genes by individual, grouped by cohort.

### CAZymes are increased in captive bears

Carbohydrate-active enzymes (CAZymes) enrichment suggests that the microbial community is well-equipped to utilize complex carbohydrates, such as plant fiber (Bohra et al. 2019). We investigated microbiome CAZyme profiles to better understand how lifestyle may influence energy metabolism. Captive bears have significantly increased CAZymes compared to wild bears (p = 0.02) (Fig. 5a). This contradicts our initial hypothesis that CAZymes would be highest in wild bears, which consume a vegetation-rich diet, however, our wild bear samples were collected in August when wild bears are preferably eating berries, seeds, and nuts. In contrast, captive bears are fed more standardized diets with much less seasonal fluctuation (Tables S2-5). Habituated bears had similar CAZyme abundances to wild bears, which could be due to the consumption of high fiber anthropogenic foods, such as garden plants, produce, and processed foods. Alternatively, it may indicate that habituated bear diets are supplemented with natural vegetation. No specific CAZyme classes were significantly increased in proportion between the cohorts (Fig. 5b). These findings highlight the interplay between microbiome energy metabolism and anthropogenic versus natural-sourced nutrients.

**Figure 5.**
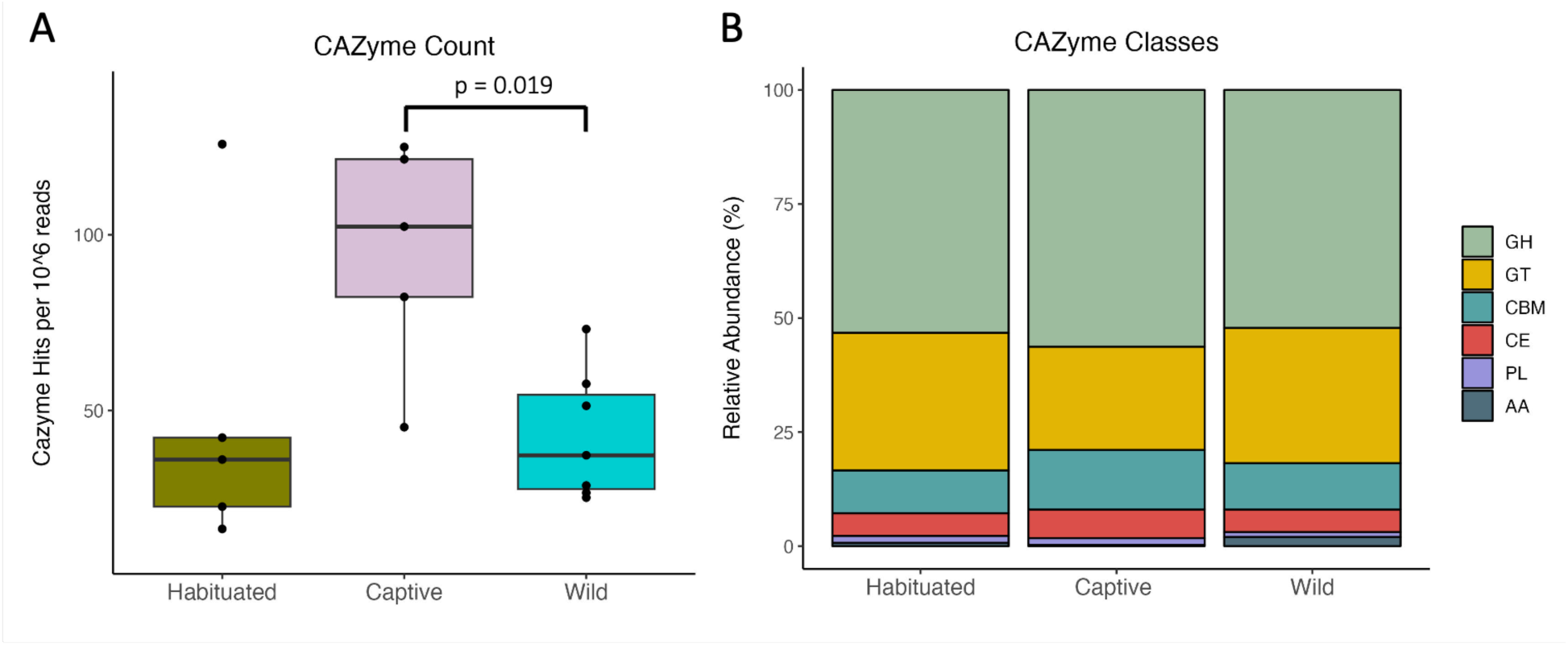
CAZyme profiles. A. CAZyme count normalized by read depth. B. CAZyme class abundances by cohort: GH = Glycoside Hydrolases, GT = Glycosyltransferases, CBM = Carbohydrate-Binding Modules, CE = Carbohydrate Esterases, PL = Polysaccharide Lyases, AA = Auxiliary Activities

## DISCUSSION

Wildlife habituation is a threat to human and animal safety, but the physiological ramifications of habituation are poorly understood. Here, we use the gut microbiome as a non-invasive proxy to interrogate how habituation, conditioning to human foods, and captivity affect wildlife health. We sampled three black bear lifestyle cohorts: wild bears with minimal human contact, captive bears with standardized diets and access to medical intervention, and habituated bears with reduced fear of humans who actively seek out human-sourced nutrients. Using amplicon and shotgun sequencing, we find a strong association between bear lifestyle and microbiome composition and function. These findings demonstrate that habituation is not only detrimental due to property damage, increased boldness, and lethal removal, but also has tangible impacts on animal and environmental health.

Our study design allowed us to directly relate cohorts of anthropogenic exposure to microbiome features in black bears. Though quite ecologically divergent, all bear species have distinctive short intestines and short, undeveloped colons that contribute to their microbiome similarities (Stevens; Schwab et al.). Despite being of the order Carnivora, wild black bears are omnivores whose diets are mostly vegetation (Hopkins et al.). We found all cohorts to be dominated by Firmicutes with high levels of Proteobacteria. This finding corroborates general microbiome compositional trends in mammals (Ley et al.) as well as previous 16S rRNA analyses done on various bear species (Sierra J. Gillman et al.; C. Song et al.; Sommer et al.; Jin et al.; Zhu et al.; Watson et al.). Broadly, we observe taxonomic differences between our bear groups that were likely driven by diet. Compared with wild bears, captive bears have enriched Peptostreptococcaceae, which is a common marker in more carnivorous carnivore microbiomes (Heni et al.; S. J. Song et al.), likely due to the reliable consumption of meat in captive bears.

Enterobacteriaceae, enriched in habituated bears compared to wild, is associated with inflammation (Moreira De Gouveia et al.) and has also been reported to disturb cellular pathways accelerating aging (Boopathi et al.). Interestingly, there is no significant difference in alpha diversity between the cohorts, which contradicts several studies that have found lower diversity in captive mammals (Kohl et al., McKenzie et al.). This finding may be driven partially by the exclusion of fungi in 16S rRNA data, which are enriched in wild bears (Supp. Results) and likely sourced from soil and plant matter. Overall, our findings demonstrate that lifestyle differences are sufficient to alter microbiome composition.

Next, we leveraged whole-metagenome shotgun sequencing to specifically interrogate microbiome features with functional links to host health. First, we focused on profiling antimicrobial resistance genes. Wildlife microbiomes have emerged as an important and understudied reservoir for antimicrobial resistance and have been shown to harbor AMR of clinical importance (Hassell et al.; Skarżyńska et al.; Swift et al.; Lagerstrom and Hadly; Argudín et al.). AMR genes are naturally coded for by microbial species and can also be acquired through chronic exposure to antimicrobial drugs as microbes evolve and select for resistance mechanisms (WHO, 2024). Much of the AMR studies in wildlife microbiomes have been limited in scope, either due to a specific focus on Enterobacteriaceae, particularly *Escherichia coli* (Carroll et al.; Chong et al.; Hassell et al.; Lagerstrom and Hadly; Radhouani et al.; Swift et al.), or due to the low throughput of culture-based approaches. Short-read sequencing allowed us to comprehensively interrogate AMR across the microbiome at the gene level.

We expected AMR levels to be highest in captive bear microbiomes, a pattern that has been observed in other animals (Bornbusch and Drea; Huang et al.) and is attributed to medical treatment and consumption of foods treated with antimicrobials such as crops and meat.

However, our results show habituated bears have significantly higher levels of AMR than both wild and captive bears. This suggests there is substantial exposure to antimicrobials in habituated bears, likely via human-sourced foods and waste (Skarżyńska et al.; Wang et al.). In primates, higher levels of AMR have been reported in individuals living in anthropogenic landscapes when compared to their counterparts experiencing less human-contact (Bornbusch and Drea; Chong et al.). Our findings warrant further investigation into AMR in wildlife, especially those residing in urban and peri-urban areas. AMR genes spread through the environment via agricultural runoff, wastewater discharge, and animal feces, and can persist and proliferate (Arnold et al.; Skandalis et al.). This environmental dissemination contributes to the global emergence of multidrug-resistant pathogens, limiting the effectiveness of antimicrobials crucial for treating bacterial infections in humans and animals (Booton et al.; Radhouani et al.). In particular, as urbanization encroaches upon wildlife habitat, human-wildlife and livestock-wildlife interactions increase and the potential for crossover increases (S. Lee et al.; Peng et al.). These findings stand as an additional caution towards the overuse and environmental dissemination of antimicrobials.

Environmental toxins impact wildlife in lethal as well as sublethal ways, such as causing disease, inducing inflammation, and altering gut microbiome composition (Nalage et al.). The reverse is true as well: toxins produced by wildlife gut microbiota are excreted via feces and disseminated into the environment (Fu et al.). Toxin profiling of gut microbiomes most commonly targets specific toxins of interest (Hou et al.). To our knowledge this is the first study to perform untargeted toxin profiling of wildlife microbiota. Toxin abundance was highest in captive bears, and is largely attributable to a high abundance of enterotoxins. Enterotoxins disrupt the normal functioning of the gut epithelium, often resulting in diarrhea and conditions such as gastroenteritis (Zhang et al.). Captivity is typically characterized by limited enclosures, regulated diet, decreased interactions with other individuals and species, medical treatment, increased human contact, and sterile environments when compared to natural habitats (Rabin). Captivity has been shown to influence behavior, stress, microbiome composition, and even fitness of animals -though these findings are highly species-specific (Farquharson et al.; Fischer and Romero; McKenzie et al.; Rabin). Bears at captive facilities may be predisposed to health problems such as gastrointestinal distress because they are rescued individuals being rehabilitated. While we acknowledge the importance of captivity for wildlife management, rehabilitation and conservation, our findings suggest further complications of captivity.

Carbohydrate-active enzymes (CAZymes) enable microbiota to efficiently metabolize plant polymers and fibers that are otherwise indigestible to the host (Wardman et al.). The resulting production of short-chain fatty acids essential to metabolism, immune function, and gut physiology (Koh et al.) contribute to the resilience and adaptability of wildlife gut microbiomes. In general, documented wildlife CAZyme profiles cluster based on dietary classification (herbivore, omnivore, and carnivore) rather than species phylogeny (Muegge et al.). Increased dietary fiber has been associated with higher CAZyme abundance (Bhattacharya et al.). Thus, we expected CAZyme abundance to be the highest in wild bears, which we assumed would be consuming the highest dietary proportion of vegetation. Instead, we observed significantly higher CAZyme abundance in captive bears, suggesting captive bears are consuming the highest proportion of complex plant fibers. This result may be attributable to seasonal cycling in wild bear diets.

Samples were collected in the late summer, when wild bears are preferably eating berries, seeds, and nuts rather than high fiber foods. Future work could characterize CAZyme profiles across seasons in wild bears to better distinguish between the effects of diet and broader lifestyle on microbial nutrient metabolism.

The work presented here yields important insights to the physiological and environmental repercussions of captivity and habituation. However, we acknowledge there are several limiting factors in our study design and analyses. Bears are highly seasonal animals, changing their diet, behavior, and ecological interactions across seasons (Mazur), which have been shown to correspond to microbiome composition and functional shifts (Sommer et al.). Habituation alters seasonal shifts since human-sourced nutrients such as trash are readily available year-round. Seasonal shifts should also be less dramatic in captive bears, who receive year-round food. To minimize seasonal bias, we chose wild and habituated samples for short-read sequencing that were all collected in the same month. Thus, the findings presented here represent a snapshot of a bear’s microbiome composition during their active season, but this composition may vary seasonally. Similarly, the geographic location of wild bears determines the food they have access to throughout the year. Our sample collection was opportunistic and spans several hundred miles through landscapes that have differing climates and species assemblages. We acknowledge closer geographically paired sampling of wild and habituated individuals would be ideal. We also did not account for the ages of bears, which also influences microbiome composition (Ghosh et al.; Sierra J. Gillman et al.). Finally, as a technical limitation, sequencing data analyses rely on comprehensive reference databases for taxonomic and functional assignment. The vast majority of microbiome WGS is done on human samples and therefore wildlife microbiomes harbor significant amounts of unclassified microbes and functions (Avila Santos et al.; Levin et al.; Abdill et al.). As wildlife microbiome WGS becomes more widespread, increased comprehensiveness of reference databases will enable deeper investigation into the role of previously undescribed taxa.

Wildlife habituation is dangerous for humans and wildlife and is ever-increasing as urbanization and habitat fragmentation spread globally. Here, we show that lifestyles across a human-exposure gradient drive differences in black bear gut microbiome composition and function.

Habituated bears exhibited significantly higher AMR loads, likely due to exposure to human-sourced nutrients, highlighting the need for further investigation into AMR transmission in urban wildlife. Meanwhile, captive bears exhibited the highest toxin abundance, primarily due to enterotoxins, suggesting that captivity may influence gut microbiome composition and predispose individuals to gastrointestinal distress. Captive bears also exhibited the highest CAZyme abundance, suggesting a diet richer in complex plant fibers than wild bears, likely due to seasonal shifts in wild bear foraging that emphasize lower-fiber foods like berries, seeds, and nuts in late summer. This work establishes a foundation for future studies to use the microbiome as a non-invasive proxy for describing the relationships between wildlife lifestyle and health, as well as their contributions to One Health.

