## Supplemental Material for "Habituation drives elevated antimicrobial resistance, functional, and compositional shifts in the black bear gut microbiome"

### Supplementary Figures

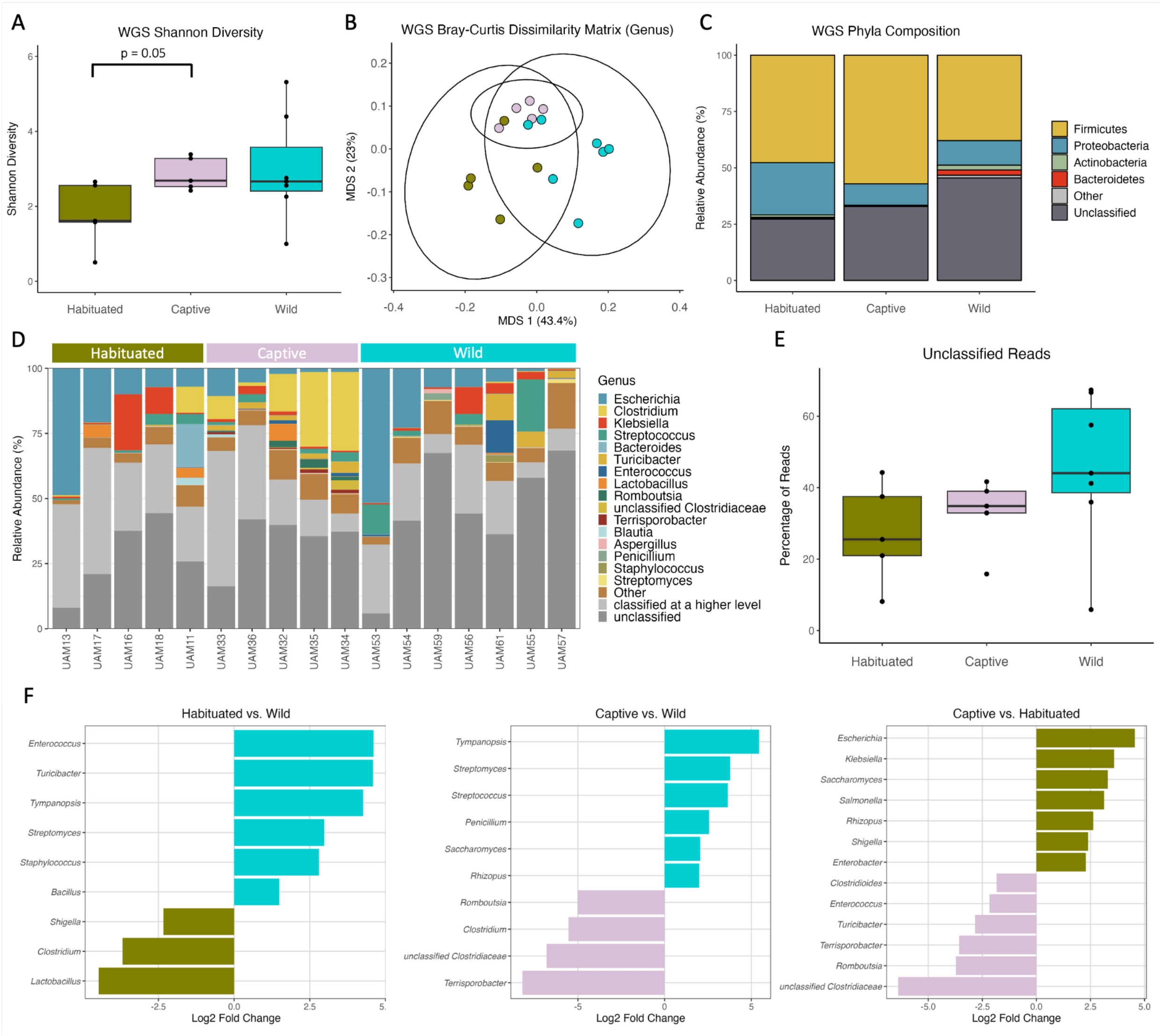


Supp. Fig 1. WGS composition at the genus level. A. Shannon diversity is not significantly different between cohorts. B. Bray-Curtis dissimilarity matrix colored by cohort. C. Phyla abundance by cohort. D. Genus relative abundance by individual, grouped by cohort. E. Percentage of unclassified reads by cohort. F. Pairwise significantly different bacterial genera.

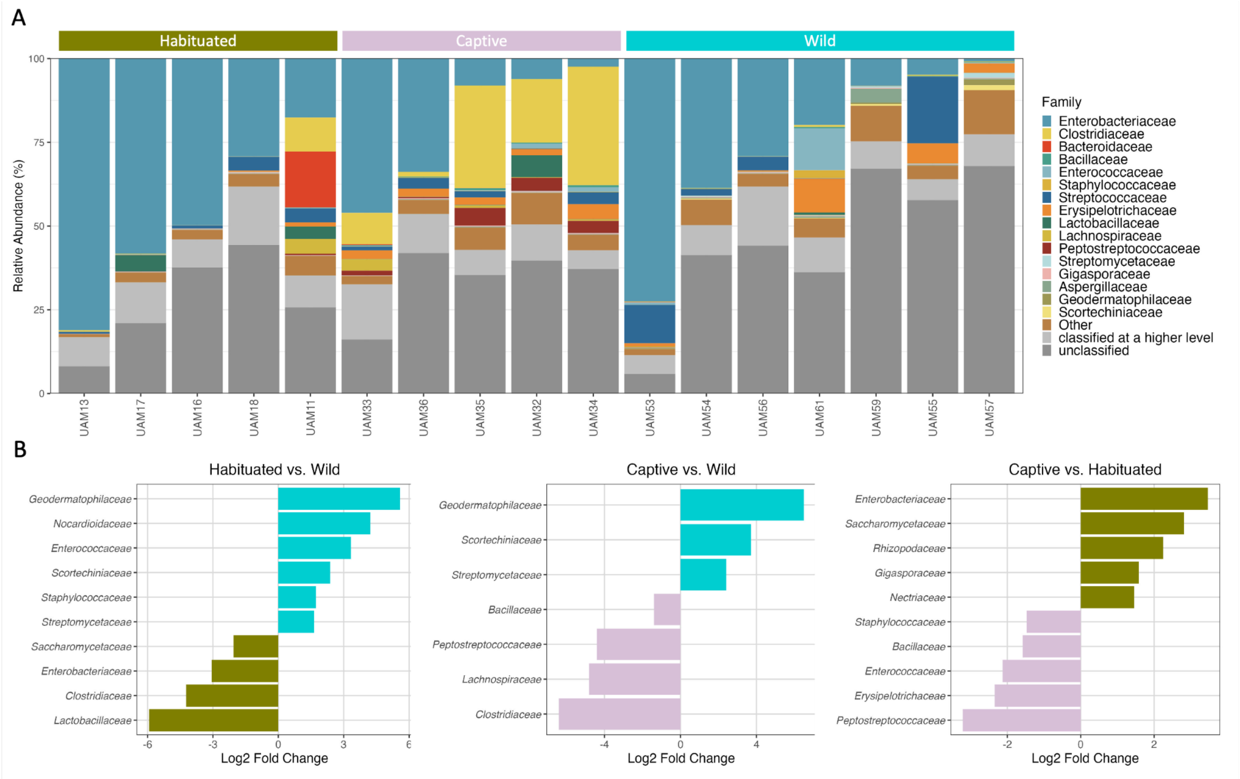


Supp. Fig 2. WGS composition at the family level. A. Family relative abundance by individual, grouped by cohort. B. Pairwise significantly different bacterial families.

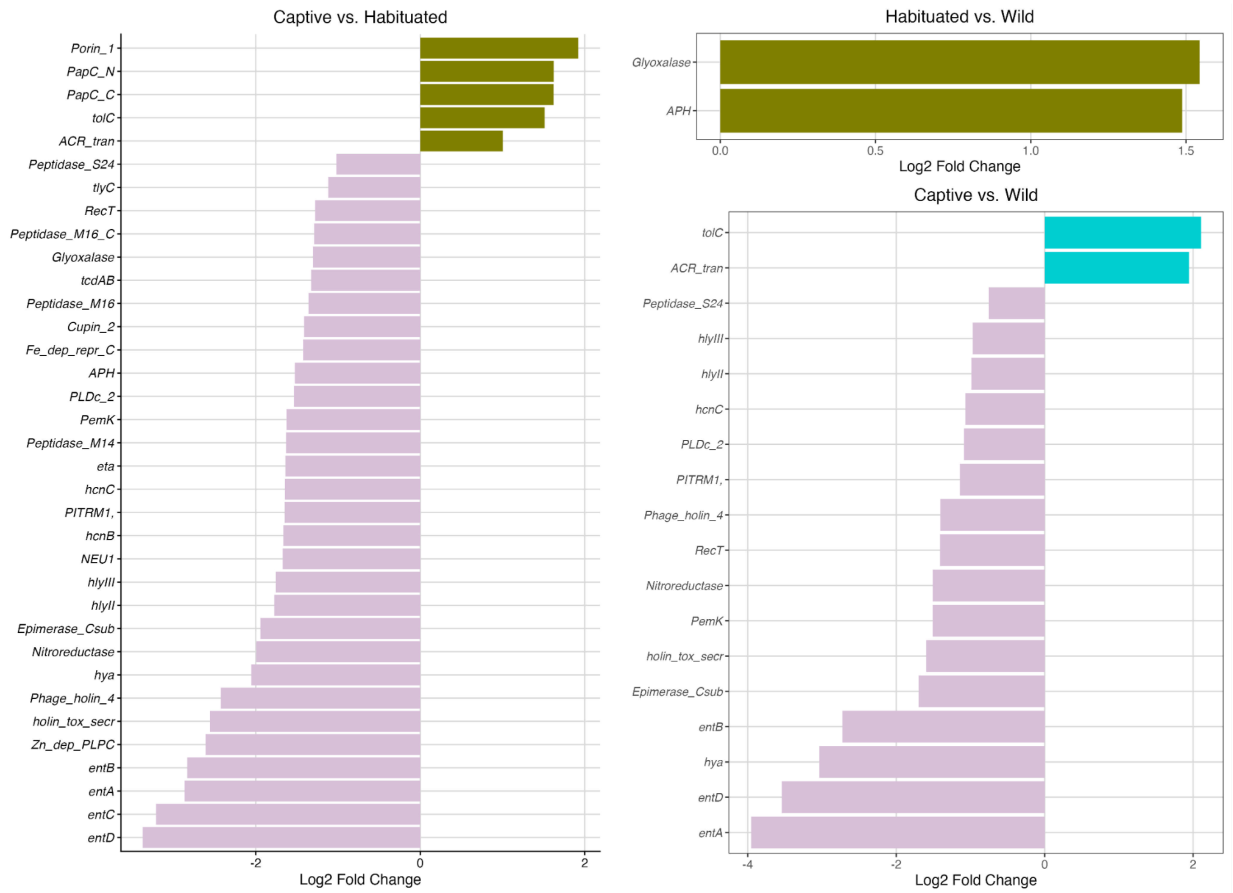


Supp Fig 3. Pairwise differential abundances of toxin-associated genes

### Supplemental Tables

**Supp. Table 1. Sample details, collection site, and date of collection**

| Study Group | County | Sample | Date of Sample Collection | Location Notes |
|---|---|---|---|---|
| Captive | Plumas | UAM003 | 5/15/2017 | WIL |
| Captive | Placer | UAM027 | 1/12/2017 | Folsom City Zoo Sanctuary |
| Captive | Placer | UAM028 | 8/30/2017 | Folsom City Zoo Sanctuary |
| Captive | Placer | UAM029 | 8/30/2017 | Folsom City Zoo Sanctuary |
| Captive | Placer | UAM030 | 8/30/2017 | Folsom City Zoo Sanctuary |
| Captive | Placer | UAM031 | 8/30/2017 | Folsom City Zoo Sanctuary |
| Captive | Sacramento | UAM032 | 10/7/2018 | PAWS (Galt) |
| Captive | Sacramento | UAM033 | 10/7/2018 | PAWS (Galt) |

| | | | | |
|---|---|---|---|---|
| Captive | Calaveras | UAM034 | 10/15/2018 | PAWS (San Andreas) |
| Captive | Calaveras | UAM035 | 10/15/2018 | PAWS (San Andreas) |
| Captive | Calaveras | UAM036 | 10/26/2018 | PAWS (San Andreas) |
| Captive | El Dorado | UAM037 | 10/17/2018 | Lake Tahoe Wildlife Care |
| Captive | El Dorado | UAM038 | 10/17/2018 | Lake Tahoe Wildlife Care |
| Captive | El Dorado | UAM039 | 11/14/2018 | Lake Tahoe Wildlife Care |
| Captive | Alameda | UAM040 | 9/20/2018 | Oakland City Zoo |
| Captive | Alameda | UAM041 | 9/20/2018 | Oakland City Zoo |
| Captive | Alameda | UAM042 | 9/20/2018 | Oakland City Zoo |
| Captive | Alameda | UAM043 | 9/20/2018 | Oakland City Zoo |
| Captive | Alameda | UAM044 | 9/20/2018 | Oakland City Zoo |
| Captive | Alameda | UAM045 | 9/20/2018 | Oakland City Zoo |
| Captive | Alameda | UAM046 | 9/20/2018 | Oakland City Zoo |
| Captive | Alameda | UAM047 | 9/20/2018 | Oakland City Zoo |
| Wildland | Mono | UAM020 | 7/17/2017 | Glass Mountains |
| Wildland | Mono | UAM021 | 8/3/2017 | |
| Wildland | Mono | UAM022 | 7/9/2017 | |
| Wildland | Mono | UAM023 | 7/31/2017 | |
| Wildland | Mono | UAM024 | 7/11/2017 | |
| Wildland | Mono | UAM025 | 7/27/2017 | |
| Wildland | El Dorado | UAM048 | 11/11/2018 | Ice House |
| Wildland | El Dorado | UAM049 | 11/11/2018 | Ice House |
| Wildland | Mono | UAM050 | 6/29/2017 | |
| Wildland | Mono | UAM051 | 6/2/2017 | |
| Wildland | San Bernardino | UAM052 | 10/15/2018 | Crystal Lake |
| Wildland | San Bernardino | UAM053 | 8/23/2018 | Mission Creek |
| Wildland | San Bernardino | UAM054 | 8/13/2018 | North Lake Arrowhead |
| Wildland | San Bernardino | UAM055 | 8/14/2018 | Oak Glenn Conservation Camp |
| Wildland | Los Angeles | UAM056 | 8/16/2018 | Bear Paw (Angeles National Forest) |
| Wildland | Los Angeles | UAM057 | 8/30/2018 | Bear Paw (Angeles National Forest) |

| | | | | |
|---|---|---|---|---|
| Wildland | Los Angeles | UAM058 | 11/17/2018 | Angeles National Forest |
| Wildland | Los Angeles | UAM059 | 9/14/2018 | Ice House Canyon |
| Wildland | Los Angeles | UAM060 | 11/28/2018 | Angeles National Forest |
| Wildland | Los Angeles | UAM061 | 8/15/2018 | Angeles National Forest |
| Wildland | Los Angeles | UAM062 | 11/27/2018 | Angeles National Forest |
| Wildland | Los Angeles | UAM063 | 11/27/2018 | Angeles National Forest |
| Urban-wildland | Tehama | UAM001 | 4/6/2017 | Wildlife Investigations Lab |
| Urban-wildland | Unknown | UAM002 | 5/5/2017 | |
| Urban-wildland | Washoe | UAM007 | 8/8/2017 | Incline Village, NV |
| Urban-wildland | Washoe | UAM008 | 6/14/2017 | Incline Village, NV |
| Urban-wildland | Washoe | UAM009 | 6/27/2017 | Reno, NV |
| Urban-wildland | Douglas | UAM011 | 8/29/2017 | Zephyr Cove, NV |
| Urban-wildland | Douglas | UAM013 | 7/28/2017 | Gardnerville, NV |
| Urban-wildland | Washoe | UAM014 | 7/8/2017 | Sand Harbor, NV |
| Urban-wildland | Washoe | UAM016 | 8/4/2017 | Washoe Valley |
| Urban-wildland | Washoe | UAM017 | 8/4/2017 | Washoe Valley |
| Urban-wildland | Douglas | UAM018 | 8/20/2017 | Zephyr Cove |
| Urban-wildland | Douglas | UAM019 | 6/8/2017 | Zephyr Cove |
| Urban-wildland | Douglas | UAM064 | 10/27/2018 | Gardnerville, NV |
| Urban-wildland | N/A | UAM065 | 11/3/2018 | Carson City, NV |
| Urban-wildland | Douglas | UAM066 | 8/7/2018 | Stateline, NV |
| Urban-wildland | Storey | UAM067 | 7/13/2018 | Virginia City Highlands, NV |
| Urban-wildland | Washoe | UAM068 | 7/9/2018 | Incline Village, NV |
| Urban-wildland | Storey | UAM069 | 6/4/2018 | Virginia City Highlands, NV |

| | | | | |
|---|---|---|---|---|
| Urban-wildland | Douglas | UAM070 | 6/12/2018 | Gardnerville, NV |
| Urban-wildland | Douglas | UAM071 | 9/29/2018 | |
| Urban-wildland | Douglas | UAM072 | 10/13/2018 | Gardnerville, NV |

**Supp. Table 2. Folsom City Zoo Sanctuary bear diets**

| Produce | Bear 1 | Bear 2 | Bear 3 | Bear 4 | Bear 5 |
|---|---|---|---|---|---|
| Lettuce (head) | 2 | 2 | 2 | 2 | 2 |
| Dandelion greens (bunch) | 1 | 1 | 1 | 1 | 1 |
| Berries (box) | 1 | 1 | 1 | 1 | 1 |
| Grapes (bag) | 1/2 | 1/2 | 1/2 | 1/4 | 1/4 |
| Apples | 2 | 2 | 2 | 2 | 2 |
| Oranges | 2 | 2 | 2 | 2 | 2 |
| Yams | 1 | 1 | 1 | 1 | 1 |
| Corn | 1 | 1 | 1 | 1 | 1 |
| Carrots | 2 | 2 | 2 | 2 | 2 |
| Artichokes | 1 | 1 | 1 | 1 | 1 |
| Celery (bunch) | 1 | 1 | 1 | 1/2 | 1/2 |
| Beets | 1 | 1 | 1 | 1 | 1 |
| Pineapple/melon | 1/4 | 1/4 | 1/4 | 1/4 | 1/4 |
| Pears | 2 | 2 | 2 | 2 | 2 |
| Canidae dog kibble (cup) | 1 | 1 | 1 | 1 | 1 |
| Varied raw meat (no fish) 1x/week (pounds) | 1 | 1 | 1 | 1 | 1 |
| Supplements | Glycoflex | glycoflex | glycoflex | glycoflex | glycoflex |
| | omega 3/6 | omega 3/6 | omega 3/6 | omega 3/6 | omega 3/6 |
| | | | meloxicam | | |
| | | | adequan | | |

**Supp. Table 3. Lake Tahoe Wildlife Care bear diets**

| Produce | Bear 1 | Bear 2 | Bear 3 |
|---|---|---|---|
| Apples (pounds) | 7 | 7 | 7 |
| Grapes (bag) | 2 | 2 | 2 |
| Peaches or pears (pieces) | 5 | 5 | 5 |
| Raspberries/blackberries (carton) | 2 | 2 | 2 |
| Watermelon (seasonally) | 1/2 | 1/2 | 1/2 |
| Lettuce (head, when available) | 2 | 2 | 2 |
| Trout; kokanee or mackinaw | 1 fish | 1 fish | 1 fish |
| Deer meat (pounds, when available) | 2 | 2 | 2 |
| Mealworms (pounds) | 1/16 | 1/16 | 1/16 |
| Mazuri bear chow (pounds) | 1 | 1 | 1 |

**Supp. Table 4. Oakland Zoo bear diets**

| Produce | Bear 1 | Bear 2 | Bear 3 | Bear 4 |
|---|---|---|---|---|
| Mixed greens* (pounds) | 5 1/2 | 5 1/2 | 5 1/2 | 5 1/2 |
| Root vegetables** (pounds) | 1 | 1 | 1 | 1 |
| Variety vegetables*** (pounds) | 3 | 3 | 3 | 3 |
| Variety fruits**** (pounds) | 1 | 1 | 1 | 1 |
| Mixed raw nuts (pounds) | 1/4 | 1/4 | 1/4 | 1/4 |
| Mealworms (pounds) | 1/12 | 1/12 | 1/12 | 1/12 |
| Mazuri kibble (Omnivore diet) pounds | 1.5 | 1.5 | 1.5 | 1.5 |
| Mazurki kibble (Bear maintenance) pounds | 1 | 1 | 1 | 1 |

| | | | | |
|---|---|---|---|---|
| Crickets and cockroaches (1x/week) pounds | 1/4 | 1/12 | 1/12 | 1/12 |
| Raw turkey (1x/week) pounds | 2-3 | 2-3 | 2-3 | 2-3 |
| Rabbit (1x/week) whole | 1 | 1 | 1 | 1 |
| Pork neck bones (1x/week) pounds | 1-2 | 1-2 | 1-2 | 1-2 |
| Cow femur bones (1x/week) pounds | 1 ½-2 | 1 ½-2 | 1 ½-2 | 1 ½-2 |
| Raw horse loin (1x/week) pounds | 1 2/3 | 1 2/3 | 1 2/3 | 1 2/3 |
| Raw chicken (pounds) | 2-4 | 2-4 | 2-4 | 2-4 |

**Mixed greens may include but are not limited to iceberg lettuce, romaine lettuce, spinach, and cabbage*

***Root vegetables may include but are not limited to yams, sweet potatoes, carrots, celery root, beets, and turnips*

****Variety vegetables may include but are not limited to pumpkin, celery, corn, peas, cucumber, tomato, avocado, jicama, and radish*

*****Variety fruits may include but are not limited to melons, oranges, papaya, grapes, plums, mango, pear, berries, nectarines, peaches*

**Supp. Table 5. PAWS bear diets**

| Produce | Bear 1 (Galt) | Bear 2 (Galt) | Bear 3 (San Andreas) | Bear 4 (San Andreas) | Bear 5 (San Andreas) | Bear 6 (San Andreas) |
|---|---|---|---|---|---|---|
| Lettuce (head) | 1 | 1 | 1 | 1 | 1 | 1 |
| Carrots (pieces) | 5 | 5 | 5 | 5 | 5 | 5 |
| Avocado (pieces) | 2 | 2 | 2 | 2 | 2 | 2 |
| Squash (pounds) | 4 | 4 | 4 | 4 | 4 | 4 |
| Sweet potato (pounds) | 5 | 5 | 5 | 5 | 5 | 5 |
| Black/Blueberries (cup) | 1 | 1 | 1 | 1 | 1 | 1 |
| Tomato (pieces) | 3 | 3 | 3 | 3 | 3 | 3 |
| Melon (occasionally) | 1/2 | 1/2 | 1/2 | 1/2 | 1/2 | 1/2 |
| Raw chicken quarters (times per week) | 3-5 | 3-5 | 3-5 | 3-5 | 3-5 | 3-5 |
| Fish (Atlantic salmon or tilapia, occasionally) | 1 | 1 | 1 | 1 | 1 | 1 |

| | | | | | | |
|---|---|---|---|---|---|---|
| Mazuri bear chow (cup) | 1 | 1 | 1 | 1 | 1 | 1 |
| Cosequin (grams) | 6.6 | 6.6 | 6.6 | 6.6 | 6.6 | 6.6 |
| Milk Thistle (grams) | 25 | 25 | 25 | 25 | 25 | 25 |
| Missing Link (grams) | 16 | 16 | 16 | 16 | 16 | 16 |

### SUPPLEMENTAL RESULTS

#### Short-read sequencing compositional results

In short-read sequencing, habituated bears have the lowest Shannon diversity followed by captive and then wild bears. Lower gut microbiome diversity has been associated with dysbiosis and various disease and infection states (Kriss et al.). This is the result we expected, assuming the unnatural diets of habituated bears would negatively influence microbiome function, but did not observe in our 16S rRNA sequencing data. Despite Shannon diversity being highest in the wild bear cohort, they have the least classified metagenome: the highest percentage of reads that were not classifiable to any known microbial database. In addition, the fewest number of genes were identified in the wild cohort metagenome. This is counterintuitive because with the highest Shannon diversity as well as the most 16S rRNA genes assembled, we expect to identify the most genes in wild bears. Our contrasting result is likely an artifact of strongly human-biased databases, both for microbial species and characterized genes (Avila Santos et al.). Captivity and urbanization have been shown to humanize wildlife gut microbiomes, which is why databases would better represent captive and habituated bears that more closely interact with humans and consume human-sourced nutrients (Clayton et al.; Dillard et al.). Contrary to several mammalian studies, including giant pandas and Andean spectacled bears, we did not detect significantly lower diversity in captive bears using either sequencing method (Borbón-García et al.; Guo, Mishra, Wang, et al.).

**Short-read sequencing shows reduced diversity in habituated bears**

Contrary to the 16S rRNA results, short-read sequencing of a subset of bears shows reduced Shannon diversity in habituated bears. In addition, the compositional separation of the cohorts is much more pronounced than in 16s rRNA gene sequencing, with habituated bears clustering closely together in between captive and wild cohorts. Wild bears still show the largest spread in diversity and across the Bray Curtis dissimilarity matrix. Despite assembling the most 16S rRNA genes in wild bears, they have the fewest classified reads and genes when corrected for sequencing depth. Captive, habituated, and wild bears had mean unclassified read values of 27.3%, 32.86%, and 45.52% and mean read counts of 33.7, 22.9, 34.5, million, respectively.

Bray-Curtis dissimilarity clustering of shotgun data shows much tighter cohort grouping than 16S rRNA sequencing, with habituated bears grouping closely together as an intermediate between captive and wild bears, who are more widely spread. Short-read MDS axes also explain much more variance than the 16S rRNA data at both the family and genus level. This result paired with lower microbiome diversity could mean that habituated bears consume a more standardized diet than wild bears on average. Human-sourced nutrient consumption is very opportunistic but also very reliable, meaning habituated bears do not need to accommodate seasonally changing diet items (Mazur). Diet homogeneity and reduced quality could be driving a less diverse and adaptable microbiome in habituated bears with less variation between individuals. Other physiological processes could also be influencing habituated microbiomes, such as disrupted temporal activity, pollutant and toxin exposure, and increased stress levels. Subsampling the 16S rRNA data to include only the individuals who were also short-read sequenced results in similar, though less striking, patterns of clustering (supp fig).

Short-read sequencing is a powerful tool for investigating microbiomes at the gene level. However, classification rates can suffer because whole genome sequencing on wildlife microbiomes is rare, and those studies that have been performed have shown they contain multitudes of unclassified diversity (Avila Santos et al.; Levin et al.). Abundances of species are necessarily based only on classifiable reads, which can bias the results. This problem is remedied by studies that perform deep WGS using either short- or long-read sequencing and then construct metagenomic bins to assemble novel genomes to then add to microbial databases. Deeper sequencing, including long-read sequencing, paired with metagenomic binning and genome reconstruction would help us better understand the origins and function of our unclassified data (Levin et al.).
